# Phen2Gene: Rapid Phenotype-Driven Gene Prioritization for Rare Diseases

**DOI:** 10.1101/870527

**Authors:** Mengge Zhao, James M. Havrilla, Li Fang, Ying Chen, Jacqueline Peng, Cong Liu, Chao Wu, Mahdi Sarmady, Pablo Botas, Julián Isla, Gholson Lyon, Chunhua Weng, Kai Wang

**Affiliations:** Center for Cellular and Molecular Therapeutics, Children’s Hospital of Philadelphia, Philadelphia, PA 19104, USA; Department of Bioengineering, University of Pennsylvania, Philadelphia, PA 19104, USA; Department of Biomedical Informatics, Columbia University Medical Center, New York, NY 10032, USA; Division of Genomic Diagnostics, Children’s Hospital of Philadelphia, Philadelphia, PA 19104, USA; Department of Pathology and Laboratory Medicine, University of Pennsylvania Perelman School of Medicine, Philadelphia, PA 19104, USA; Foundation 29, Pozuelo de Alarcon, Madrid, Spain; Orphan Drug Committee, European Medicine Agency, Amsterdam, The Netherlands; Institute for Basic Research in Developmental Disabilities (IBR), Staten Island, New York, USA

**Author notes:** These authors contributed equally to this work (Mengge Zhao, James M. Havrilla, Li Fang).

## Abstract

Human Phenotype Ontology (HPO) terms are increasingly used in diagnostic settings to aid in the characterization of patient phenotypes. The HPO annotation database is updated frequently and can provide detailed phenotype knowledge on various human diseases, and many HPO terms are now mapped to candidate causal genes with binary relationships. To further improve the genetic diagnosis of rare diseases, we incorporated these HPO annotations, gene-disease databases, and gene-gene databases in a probabilistic model to build a novel HPO-driven gene prioritization tool, Phen2Gene. Phen2Gene accesses a database built upon this information called the HPO2Gene Knowledgebase (H2GKB), which provides weighted and ranked gene lists for every HPO term. Phen2Gene is then able to access the H2GKB for patient-specific lists of HPO terms or PhenoPackets descriptions supported by GA4GH (http://phenopackets.org/), calculate a prioritized gene list based on a probabilistic model, and output gene-disease relationships with great accuracy. Phen2Gene outperforms existing gene prioritization tools in speed, and acts as a real-time phenotype driven gene prioritization tool to aid the clinical diagnosis of rare undiagnosed diseases. In addition to a command line tool released under the MIT license (https://github.com/WGLab/Phen2Gene), we also developed a web server and web service (https://phen2gene.wglab.org/) for running the tool via web interface or RESTful API queries. Finally, we have curated a large amount of benchmarking data for phenotype-to-gene tools involving 197 patients across 76 scientific articles and 85 patients’ de-identified HPO term data from CHOP.

## Introduction

Rapid and accurate genetic diagnosis of Mendelian diseases is necessary to optimize both treatment and management strategies and implement precision medicine. Compared to traditional single-gene tests or gene panels, recent efforts have utilized next-generation sequencing (NGS) technologies, such as whole exome sequencing (WES) and whole genome sequencing (WGS). The intent of the NGS effort is to improve diagnostic rates, enhance time efficiency and decrease overall financial burdens^1–5^. However, due to the substantially larger pool of candidate genes created by NGS data, sequence interpretation has become a major hurdle in diagnostic settings. Computational approaches that streamline the diagnostic workflow and shorten the analytical turnaround time are needed.

The Human Phenotype Ontology (HPO) database^6^ associates human diseases with phenotypic abnormalities. These terms possess ever-increasing interoperability with other ontologies^7–11^ and allow for computational deep phenotyping, making it the prevailing standard terminology for human phenotypes. We have developed a few computational tools^12–14^ that use phenotype data for gene prediction and prioritization. Although these methods are useful, they cannot provide real-time decision support in clinical settings; for example, users may need to add or remove one phenotype term for a patient and immediately observe how the candidate gene list changes.

Recent studies have shown the utility of incorporating phenotype data such as HPO terms in identifying causal genes from NGS data, which increases time efficiency and diagnostic yields^15^. There are phenotype analysis tools that use HPO terms to prioritize candidate causal genes: Phevor^16^, VarElect^17^, and OVA^18^. These HPO terms can be supplied by diagnostic labs, clinical geneticists, doctors, or natural language processing (NLP) algorithms that parse doctors’ notes such as Doc2HPO14. The prioritized list of genes can then be combined with NGS data to identify potential disease genes. The downside of these tools, however, is that they require gene lists beforehand, take longer to prioritize genes, they can only be used via web interface, and have no open source code. Thus, such tools cannot be implemented on a large scale, and cannot be integrated into existing clinical diagnostic workflows, which are often protected within institutional firewalls.

We present a new rapid, accurate, phenotype-based gene-prioritization tool called Phen2Gene. Phen2Gene takes HPO terms for a patient and generates a patient-specific ranked list of candidate genes using our precomputed database, the HPO2Gene Knowledgebase (H2GKB), in a mere second. Unlike existing binary HPO-gene annotations in the HPO annotation database, the H2GKB is a new database that links each HPO term to its own ranked list of candidate genes, each with a confidence score. These scores represent a substantial accuracy improvement over the previous version of Phenolyzer. We also provide open source code under the MIT license, together with a web server for downloading and accessing the H2GKB, and a RESTful API web service for automated queries of phenotype terms with JSON output.

## Methods

### Acquiring Patient-Specific Gene Lists with Phen2Gene

Physicians can manually curate HPO terms for their patients, which is becoming more common, or feed the patients’ notes into Doc2HPO to discover the relevant and negated HPO terms for the patient’s phenotype. Doc2HPO is a public tool that uses multiple NLP tools and algorithms to parse patient notes into HPO terms. HPO terms act as the sole form of input into Phen2Gene which then searches the H2GKB for each term’s ranked gene list.

All HPO terms under the root term ‘Phenotypic abnormality’ (HP:0000118) are recognizable by Phen2Gene. By default, Phen2Gene weights the inputted HPO terms by skewness of gene scores, as some HPO terms possess more precise information content than others^19^. Then, Phen2Gene searches the H2GKB for each input term’s gene list and sorts and ranks all the genes based on their ranks in each term’s list and the weight of each HPO term to produce a final, ranked candidate gene list (**Figure 1**).

**Figure 1.**
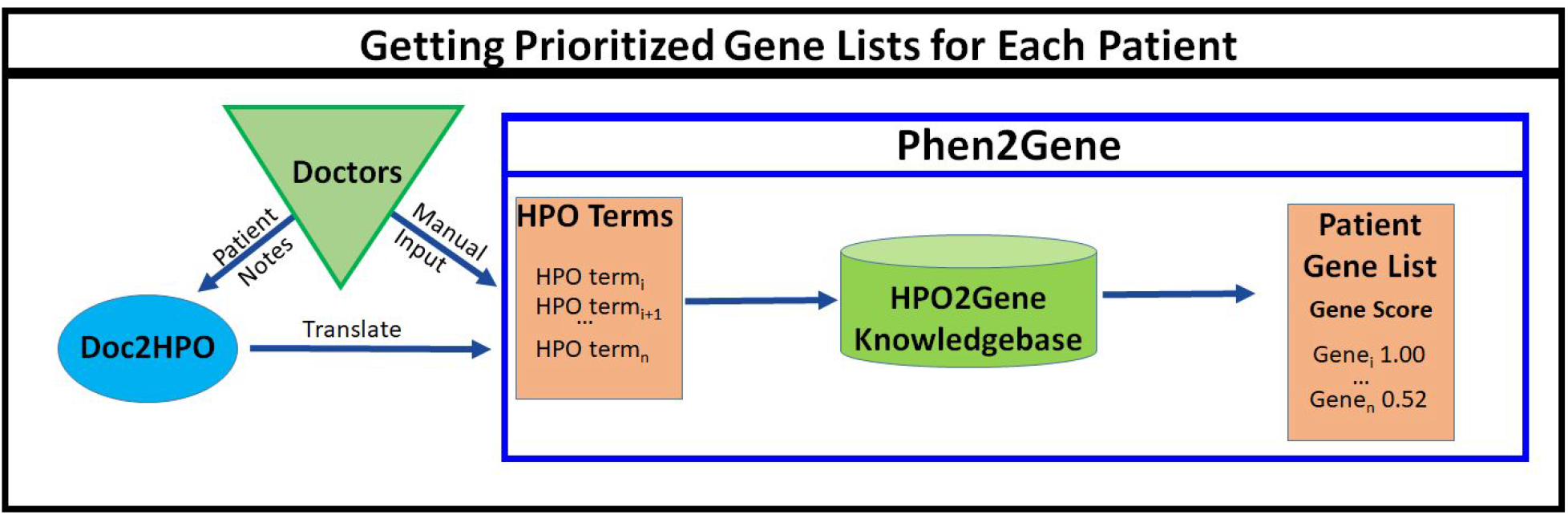
How to use Phen2Gene. Physicians or clinical geneticists can curate HPO terms themselves or provide patient notes to Doc2HPO to generate HPO terms in semi-automated fashion, and these terms will help create a candidate disease gene list using Phen2Gene.

### HPO2Gene Knowledgebase Construction

In order to construct the H2GKB, we first extract every term from the Human Phenotype Ontology Database, underneath the root term ‘Phenotypic abnormality’ (HP:0000118) (**Figure 2a**). For each HPO term, we run an enhanced version of Phenolyzer (ver. 0.4.0), dubbed Enhanced Phenolyzer, which incorporates HPO-gene annotations from the Jackson Laboratory^6^, and gene-disease annotations from OMIM^20^, ClinVar^21^, Orphanet^22^, and GeneReviews^23^. It then adds information from gene-gene databases HPRD^24^, NCBI’s Biosystems Database^25^, HGNC Gene Family^26^, and HTRI^27^, and prioritizes and outputs the associated genes.

**Figure 2.**
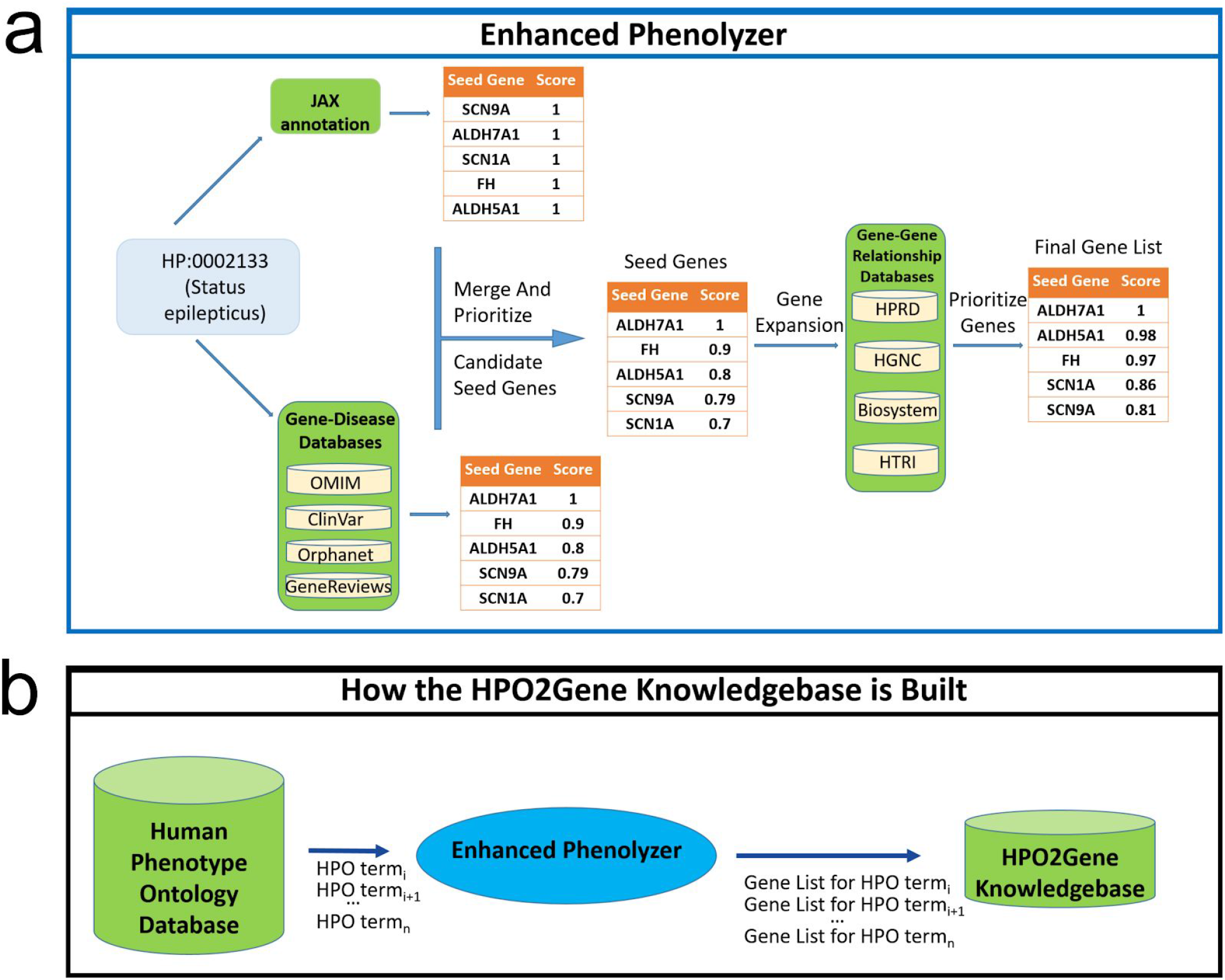
The construction of the HPO2Gene Knowledgebase. HPO terms are extracted one by one from the HPO database and passed into an enhanced version of Phenolyzer (dubbed Enhanced Phenolyzer) to create a database of ranked gene lists for all HPO terms. (a) The workflow of Enhanced Phenolyzer. (b) Construction of the HPO2Gene Knowledgebase.

This generates a ranked list of candidate causal genes for each HPO term, which are then consolidated into the H2GKB (**Figure 2b**). This precomputed H2GKB can then be rapidly accessed by Phen2Gene and used to rank lists of genes for individual patients. The H2GKB is also freely available online and downloadable from the Phen2Gene web server.

### Enhanced Phenolyzer

The original version of Phenolyzer (ver. 0.2.2) processed free-text terms supplied by users. We updated the databases inside Phenolyzer, incorporated new HPO-gene annotations from the Jackson Laboratory database, fixed some bugs, and released it as Enhanced Phenolyzer (ver. 0.4.0). Enhanced Phenolyzer contains a new function to turn HPO terms into a list of genes, by generating a prioritized gene list for each HPO term. Unlike the original Phenolyzer, Enhanced Phenolyzer first generates two seed gene sets. Seed Gene Set 1 is built on HPO-gene annotation files downloaded from the Jackson Laboratory for Genomic Medicine available at https://hpo.jax.org/app/download/annotation, while Seed Gene Set 2 construction follows the method outlined in the original Phenolyzer paper, which translates phenotype terms to disease names, and incorporates the five precompiled gene-disease databases to search for seed genes.

### Candidate Genes Prioritization

For seed genes in Set 1, we gave an equal score to each gene and HPO term pair,

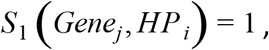

since JAX annotation only lists genes for each individual HPO term, but without quantitative scores representing the strength of associations.

In Set 2, we followed the calculation method in the original Phenolyzer for each seed gene using gene-disease databases associated with each individual HPO term, and noted as

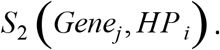

We sum up the two scores,

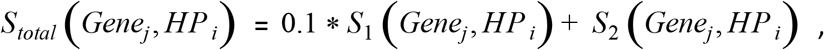

and normalized it to a range between 0 and 1 as the final seed gene score,

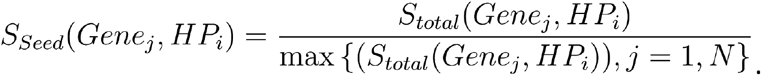

In the following steps, we used the original Phenolyzer’s method for expanding the list of candidate genes and reprioritizing the seed gene list using gene-gene databases. Then we generated the HPO2Gene-KB with Enhanced Phenolyzer.

### Weighting by Skewness

We calculated the skewness value for the distribution of all gene scores for each HPO term, and used it multiplicatively to adjust the weights of HPO terms individually. The gene score distributions vary widely from term to term. The gene score distributions of “Seizures” (HP:0001250) and “Cleft palate” (HP:0000175) demonstrate the difference in the specificity of HPO terms (**Supplementary Figure 1**). “Cleft palate” has a positively skewed gene score distribution compared to “Seizures.” For “Cleft palate,” most genes have a near zero raw score value, but for seizures the mean and standard deviation are much larger. We assume that the more skewed the gene score distribution, the greater the difference between high and low ranking genes. This discrepancy provides HPO terms with better information for their associated genes. Thus, we used Pearson’s moment coefficient of skewness to represent the skew and weight HPO terms’ gene weights:

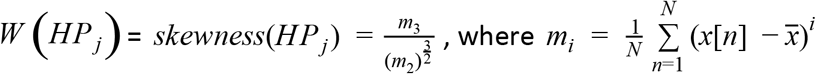

and where *skewness*(*HP_j_*) is the skewness of the gene-score distribution of *HP_j_*, which we calculated with Python 3.8 and the SciPy 1.3.1 stats module. We also created alternative weighting schemes involving no weight or informational content (**Supplemental Methods**).

### Gene Score Computation with Weighted HPO Terms

Given a set of HPO terms, *TermSet* = **{***HP_j_***}**, each HPO term is assigned a weight representing the granularity of phenotypic information given by the HPO term. In each *HP_j_*’s candidate gene list, every candidate gene has a score calculated by Enhanced Phenolyzer. It is a quantitative representation of how *gene_i_* is associated with *HP_j_*. Phen2Gene gives a weighted score to *gene_i_*, if *gene_i_* is in *HP_j_*’s candidate gene list,

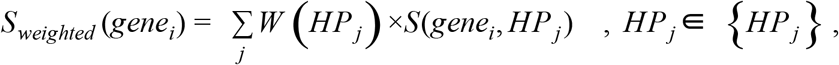

where *W*(*HP_j_*) is the assigned weight as illustrated in the previous section, *S*(*gene_i_, HP_j_*) is *gene_i_*’s score in *HP_j_*’s candidate gene list. *S*(*gene_i_, HP_j_*) = 0, if *gene_i_* is not a candidate gene of *HP_j_*. All of genes are sorted by their scores in descending order.

## Results

### General Use

Since the H2GKB is precomputed, the results for Phen2Gene are instant. The weight given to the HPO terms can be chosen or defined by the end user. The terms can be unweighted, weighted by ontology-based information content, or the skewness of gene scores for each HPO term, which is the recommended default. No prior gene list knowledge is required, and if a physician has no candidate genes, or if whole exome or whole genome sequencing cannot be performed for practical reasons (such as insurance reimbursement issues), it could help select a targeted sequencing gene panel to find variants causal for the phenotype. This process can be performed case-by-case on the web server, or using the Phen2Gene python script and thus can be scaled up massively to thousands of patients without prior gene knowledge, unlike competing HPO-to-gene software tools.

### Accuracy Evaluation with Collected Expert-Curated Phenotype Data

For our benchmark testing of Phen2Gene, we used 282 de-identified patients who were diagnosed as carrying single disease gene as our study subjects. Their study data were from 5 different sources but 3 were manually curated, hence we divided them into 4 groups. Group 1 only contained one disease gene: TAF1, but the other three groups contained numerous known and previously validated disease genes (**Table 1**). The phenotypes in these three groups were completely randomly chosen and the phenotypes are not necessarily related in any way.

**Table 1.** Curation of benchmark data set. Each dataset comes from different literature sources, except the third set, which comes directly from the Children’s Hospital of Philadelphia. Some HPO term sets have been curated by the Aho-Corasick algorithm embedded in Doc2HPO and others were manually curated by expert physicians.

| Set | Data curation | Where HPO terms are derived |
| --- | --- | --- |
| 1 | 14 cases, with 1 unique causal gene (TAF1), from 1 *American Journal of Human Genetics* article<sup>28</sup> | Doctor-curated HPO terms |
| 2 | 27 cases from Columbia University Medical Center, with 24 unique causal genes, from 1 article<sup>13</sup> | Manually curated HPO terms from doctor defined phenotypes |
| 3 | 85 cases from the Department of Genomic Diagnostics at Children's Hospital of Philadelphia, with 75 unique causal genes | Doctor-curated HPO terms |
| 4 | 72 cases, with 59 unique causal genes, from 61 *Cold Spring Harbor Molecular Case Study* articles<sup>29–89</sup>, and 83 cases from with 13 unique genes, from 13 *American Journal of Human Genetics* articles<sup>90–103</sup> | Aho-Corasick algorithm embedded in Doc2HPO with manual review removing negated and duplicated terms |

An effective way to understand how well a phenotype-based gene prioritization tool performs is to have experts curate HPO terms and phenotype information for single-gene diseases. These experts also know the causal genes for these diseases, thus aiding in assessing if a tool is able to properly rank the causal gene highly.

Each patient case in the benchmark data set has only a single causal gene that is known beforehand by the physicians who curated the patient data. This data was used to create the benchmark test between Phen2Gene and the original version of Phenolyzer (**Figure 3**).

**Figure 3.**
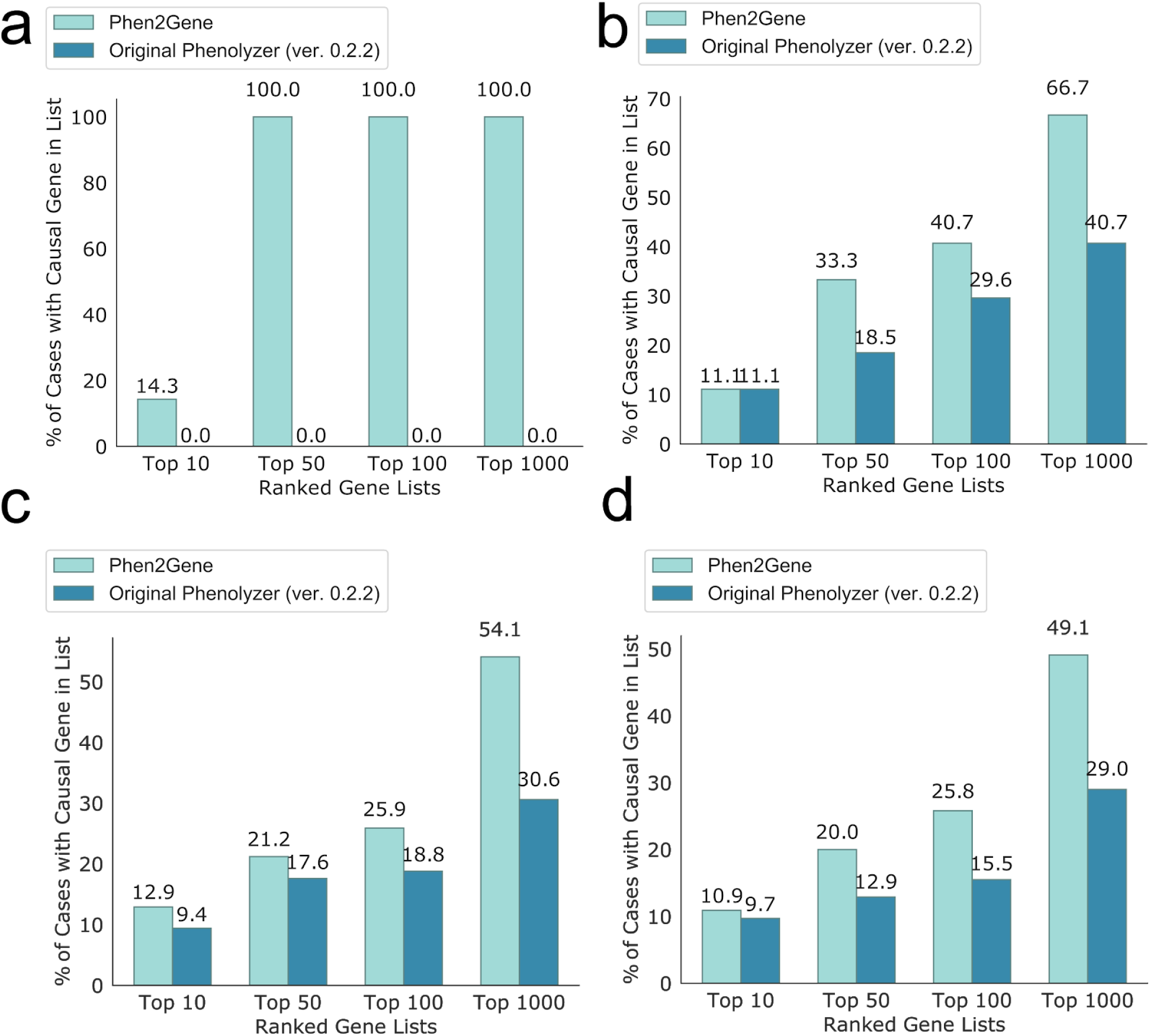
Accuracy test for Phen2Gene and the original version of Phenolyzer. The accuracy of the tool is determined by the proportion of patient cases where the causal gene was successfully identified in the top 10, top 50, top 100, and top 1000 genes for the respective tool. (a) Set 1 of patient cases for TAF1 syndrome as described in Table 1. (b) Set 2 of patient cases from Columbia University as described in Table 1. (c) Set 3 of patient cases from DGD at CHOP as described in Table 1. (d) Set 4 of patient cases from 61 CSH Mol Case Studies articles and patient cases from 13 AJHG articles as described in Table 1.

**Figure 4.**
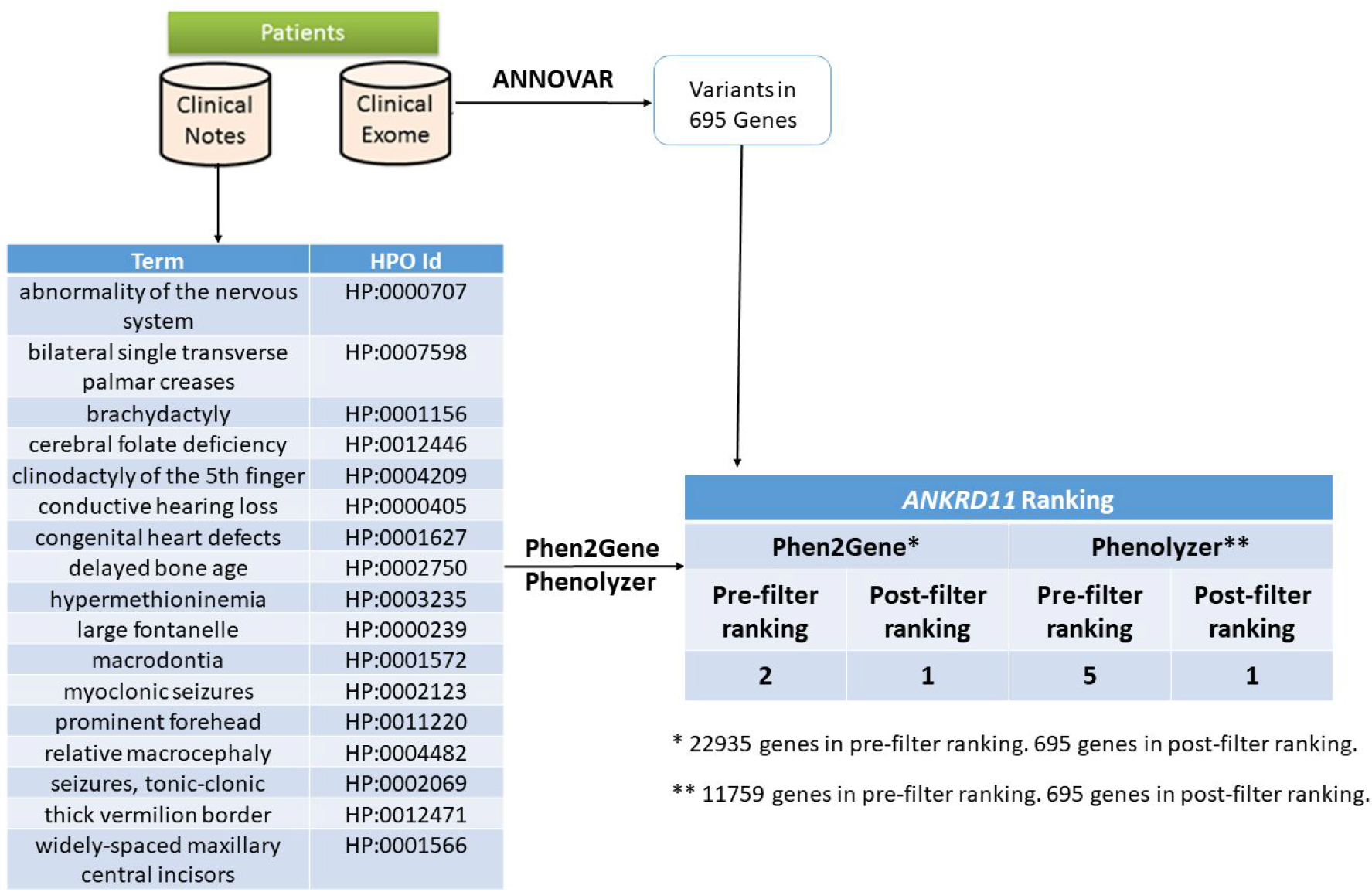
General use case. Proband has a condition with unknown genetic cause but several candidate variants annotated and filtered using ANNOVAR^104^. Clinical notes on the proband’s condition are used by Doc2HPO to generate a list of HPO terms, which act as input for Phen2Gene or Phenolyzer. These tools rank several thousand genes and by intersecting them with the candidate list of genes overlapping the variants, we obtain a list of likely candidate genes for KGB syndrome, which is known to be caused by variants in ANKRD11, shown here.

The performance of Phen2Gene and Phenolyzer varies from set to set, though overall, Phen2Gene is more accurate than Phenolyzer. The test for accuracy constitutes each tool’s ability to rank the known causal gene in the top 10, 50, 100, and 1000 genes, respectively, for each patient case, for each benchmark set. Phen2Gene represents a step forward in accuracy, and future improvements to Phenolyzer will improve the H2GKB and thus the performance of Phen2Gene even further.

Since Phen2Gene leverages the precomputed H2GKB, the speed-up in using Phen2Gene over Phenolyzer is substantial (**Table 2**). The speed with which Phen2Gene can both access and rank gene information from the H2GKB, and the fact that it can be deployed in parallel, speaks to its scalability in future large-scale phenotype analysis studies.

**Table 2.** Speed benchmark test for Phen2Gene and Phenolyzer. These represent the average, minimum and maximum runtimes of these tools in seconds and were taken from all 281 patient case runs.

| Tool | Phen2Gene | Phenolyzer |
| --- | --- | --- |
| Median Time (s) | 0.9643 | 504.54 |
| Min Time (s) | 0.5143 | 187.97 |
| Max Time (s) | 1.9264 | 1021.54 |

### General Use Case: Narrowing Down Candidate Genes For Undiagnosed Diseases

To demonstrate the real-world usage of phenotype-driven gene prioritization in clinical diagnostic settings, we performed a retrospective analysis on a previously published case. We were previously presented with a proband possessing a suspected Mendelian disease, and we performed whole exome sequencing on the proband and the parents. In a previous study, we identified a *de novo*, single-nucleotide insertion in ankyrin repeat domain 11 (ANKRD11) as the disease causal variant, and reached a genetic diagnosis of KBG syndrome, an extremely rare disease. In the current study, we evaluated whether Phen2Gene and Phenolyzer can facilitate automated gene finding from the exome data, by analyzing the proband only (i.e. without parental information).

We used the proband’s HPO terms as input for Phen2Gene and the proband’s disease and symptom terms as input for Phenolyzer. The causal gene, ANKRD11, was initially ranked 2nd and 5th by Phen2Gene and Phenolyzer, respectively, among all the genes in the genome. We intersected these gene lists with the list of candidate genes derived from genes that harbor at least one rare, protein-altering variant in the patient. ANKRD11 was ranked 1st by both Phen2Gene and Phenolyzer. This example shows how both Phen2Gene and Phenolyzer can be used to rank a causal gene in the top 10 genes based on disease and symptom information and the list of candidate genes extracted from exome sequencing. Thus human reviewers, such as clinical geneticists or genetic counselors, can review the top 10 or 50 genes and reach a genetic diagnosis with great expedition, or combine the top 1000 genes with variant information to shorten their lists of candidate genes.

## Discussion

Phen2Gene represents a change to the catalog of phenotype-to-gene software. Currently, to the best of our knowledge, no other such tool allows for scalability to thousands of patients, and often they have slow web servers that require copy-and-paste input, for one patient at a time. In addition, other tools that rank genes based on HPO terms have no open source code available — their work cannot be easily checked or improved upon by the community. In comparison, Phen2Gene is extremely fast, does not require prior gene knowledge, and does not need to be run on a web server, though the provided server is also faster than competitors’ servers. We further provide the H2GKB and the benchmark data as freely downloadable files. Compared to the annotation file that documents about 20 binary relationships between HPO terms and genes on average from Jackson Laboratory’s HPO website, the H2GKB we provide here contains weighted relationships between each HPO term and hundreds or even thousands of genes. Phen2Gene shows marked improvement over the original version of Phenolyzer, and in our future work we plan to greatly improve Phenolyzer and expand upon the H2GKB, increasing performance.

Another benefit of Phen2Gene is that it is variant agnostic. Structural variants and repeat expansions in intronic regions are known to cause disease ^105,106^, and on average, there are more than 20,000 structural variants (SVs) in the human genome^107^. Based on our calculation using the gold standard SV call set from HG002^108^ and the gene annotation file from GENCODE (v25), more than half of the structural variants overlap with genes and most overlap intronic regions. The list of tools that can score structural variants or repeat expansions is extremely small, but Phen2Gene could used to narrow down a candidate variant list containing repeat expansions or SVs.

In the future, there are several concepts which we hope to address, not the least of which is a double counting bias ubiquitous to all such HPO-to-gene tools. Some doctors’ notes may contain terms like myoclonic seizures, epilepsy and absence seizures, all of which represent three different HPO terms (HP:0002123, HP:0001250, and HP:0002121, respectively) for what is essentially the same combined condition. As a result, it may be biased towards terms mentioned more often in doctors’ notes. This redundancy can be eliminated through manual HPO term input by human experts, but is still a common issue that needs to be addressed, perhaps by downweighting similar HPO terms.

Another issue we need to handle is the issue of negated terms such as “no seizures.” Obviously, if experts input HPO terms manually, this is not a difficult issue to address, but for NLP algorithms that extract terms from doctors’ notes, we could be adding false positive HPO terms if negation is not properly detected. Conversely, using negated HPO terms to lower the ranking of negated-term-associated genes is another useful incorporation of negated term data. Integrating algorithms like DEEPEN^109^ and NegEx^110^ into tools such as Doc2HPO may help us solve this problem.

We could improve the granularity of the scoring algorithm by incorporating corpus-based information content. Phen2Gene could still be improved further, and one method for more properly assessing information content is to use HPO terms in tandem with a large body of clinical literature. This could enable us to give the proper weight to HPO terms or perhaps incorporate other terminology not covered by HPO, like UMLS, or NLP-derived classifications or clusters. There is a need for a more widely applicable terminology in the medical field, especially for diseases requiring deeper phenotyping, and this would become a useful resource for researchers doing similar work.

Finally, we can combine Phen2Gene with variant prioritization software or disease gene discovery tools such as CADD^111^, REVEL^112^, or CCR^113^, to further narrow down potential disease gene candidates. If a diagnostician has a list of genetic variants they are more likely to use one of these tools first. In the future, we hope to create a hybrid score that combines computationally derived variant scores with phenotype derived gene prioritization.

In summary, the H2GKB fills a void in the community in linking standardized phenotype terms to genes with weighted scores, and it may facilitate or inspire the development of novel computational tools that link HPO terms to genetic information, especially where whole exome/genome sequencing data is available. The Phen2Gene tool provided in this paper can rapidly access and rank this information. It has been implemented in Dx29 (http://www.dx29.ai) and Doc2HPO’s pipeline so far, and we hope to deploy it in other similar web services. Through command line tools, web servers, and RESTful API web services, we believe that Phen2Gene will greatly facilitate and expedite phenotype-driven gene prioritizations for rare diseases.

## Data Availability

The current version of Phen2Gene is 1.1.0. The source code and scripts for figures are available at https://github.com/WGLab/Phen2Gene. Additionally, we built a Phen2Gene Web Server available at http://phen2gene.wglab.org, to facilitate users who prefer to use web interface for gene prioritization. The current version of H2GKB is also downloadable at https://github.com/WGLab/Phen2Gene/releases/download/V1.1.0/H2GKB.tar.gz. All the benchmark datasets are available in the Supplementary Materials of this manuscript.

## Web Resources

HPO Website https://hpo.jax.org/app/

JAX annotations https://hpo.jax.org/app/download/annotation

Phenolyzer http://phenolyzer.wglab.org/

Doc2HPO https://impact2.dbmi.columbia.edu/doc2hpo/

Dx29 https://www.dx29.ai/

## Abbreviations

HPO: Human Phenotype Ontology
NGS: Next-generation sequencing
WES: Whole exome sequencing
WGS: Whole genome sequencing
CKD: Chronic kidney disease
NLP: natural language processing
OMIM: Online Mendelian Inheritance in Man
EHR: Electronic Health Records
NCBO: National Center for Biomedical Ontology
HG2KB: HPO2Gene Knowledgebase

## Acknowledgements

We thank the affected individuals and their family members who participated in published genetic studies to improve the diagnostic rates in clinical exome testing. We thank the original authors of the studies to provide detailed phenotype information in published manuscripts, to help us benchmark performance of different software tools. We thank the developers of the Human Phenotype Ontology for continuous development of this ontology over the past few years, which greatly facilitated and standardized clinical diagnosis of affected individuals with suspected genetic disorders. This study is supported by NIH/NLM/NHGRI grant R01LM012895 (C.W. and K.W.).

## References

1. Yang Y, Muzny DM, Reid JG, et al. Clinical whole-exome sequencing for the diagnosis of mendelian disorders. N Engl J Med. 2013;369(16):1502–1511. doi:10.1056/NEJMoa1306555

2. Eldomery MK, Coban-Akdemir Z, Harel T, et al. Lessons learned from additional research analyses of unsolved clinical exome cases. Genome Med. 2017;9(1):26. doi:10.1186/s13073-017-0412-6

3. Trujillano D, Bertoli-Avella AM, Kumar Kandaswamy K, et al. Clinical exome sequencing: results from 2819 samples reflecting 1000 families. Eur J Hum Genet. 2017;25(2):176–182. doi:10.1038/ejhg.2016.146ejhg2016146 [pii]

4. Retterer K, Juusola J, Cho MT, et al. Clinical application of whole-exome sequencing across clinical indications. Genet Med. 2016;18(7):696–704. doi:10.1038/gim.2015.148gim2015148 [pii]

5. Sawyer SL, Hartley T, Dyment DA, et al. Utility of whole-exome sequencing for those near the end of the diagnostic odyssey: time to address gaps in care. Clin Genet. 2016;89(3):275–284. doi:10.1111/cge.12654

6. Kohler S, Carmody L, Vasilevsky N, et al. Expansion of the Human Phenotype Ontology (HPO) knowledge base and resources. Nucleic Acids Res. 2019;47(D1):D1018–D1027. doi:10.1093/nar/gky1105

7. Ashburner M, Ball CA, Blake JA, et al. Gene Ontology: tool for the unification of biology. Nat Genet. 2000;25(1):25–29. doi:10.1038/75556

8. Bello SM, Shimoyama M, Mitraka E, et al. Disease Ontology: improving and unifying disease annotations across species. Dis Model Mech. 2018;11(3). doi:10.1242/dmm.032839

9. Haendel MA, Balhoff JP, Bastian FB, et al. Unification of multi-species vertebrate anatomy ontologies for comparative biology in Uberon. J Biomed Semant. 2014;5:21. doi:10.1186/2041-1480-5-21

10. Bard J, Rhee SY, Ashburner M. An ontology for cell types. Genome Biol. 2005;6(2):R21. doi:10.1186/gb-2005-6-2-r21

11. Smith CL, Goldsmith C-AW, Eppig JT. The Mammalian Phenotype Ontology as a tool for annotating, analyzing and comparing phenotypic information. Genome Biol. 2005;6(1):R7. doi:10.1186/gb-2004-6-1-r7

12. Yang H, Robinson PN, Wang K. Phenolyzer: phenotype-based prioritization of candidate genes for human diseases. Nat Methods. 2015;12(9):841–843. doi:10.1038/nmeth.3484

13. Son JH, Xie G, Yuan C, et al. Deep Phenotyping on Electronic Health Records Facilitates Genetic Diagnosis by Clinical Exomes. Am J Hum Genet. 2018;103(1):58–73. doi:S0002-9297(18)30171-X [pii] 10.1016/j.ajhg.2018.05.010

14. Liu C, Peres Kury FS, Li Z, Ta C, Wang K, Weng C. Doc2Hpo: a web application for efficient and accurate HPO concept curation. Nucleic Acids Res. 2019;47(W1):W566–W570. doi: 10.1093/nar/gkz386

15. Aerts S, Lambrechts D, Maity S, et al. Gene prioritization through genomic data fusion. Nat Biotechnol. 2006;24(5):537–544. doi:10.1038/nbt1203

16. Singleton MV, Guthery SL, Voelkerding KV, et al. Phevor combines multiple biomedical ontologies for accurate identification of disease-causing alleles in single individuals and small nuclear families. Am J Hum Genet. 2014;94(4):599–610. doi:10.1016/j.ajhg.2014.03.010 S0002-9297(14)00112-8 [pii]

17. Stelzer G, Plaschkes I, Oz-Levi D, et al. VarElect: the phenotype-based variation prioritizer of the GeneCards Suite. BMC Genomics. 2016;17(2):444. doi:10.1186/s12864-016-2722-2

18. Antanaviciute A, Watson CM, Harrison SM, et al. OVA: integrating molecular and physical phenotype data from multiple biomedical domain ontologies with variant filtering for enhanced variant prioritization. Bioinformatics. 2015;31(23):3822–3829. doi:10.1093/bioinformatics/btv473

19. Sánchez D, Batet M, Isern D. Ontology-based information content computation. Knowl-Based Syst. 2011;24(2):297–303. doi:10.1016/j.knosys.2010.10.001

20. McKusick VA. Mendelian Inheritance in Man and its online version, OMIM. Am J Hum Genet. 2007;80(4):588–604. doi:10.1086/514346

21. Landrum MJ, Lee JM, Riley GR, et al. ClinVar: public archive of relationships among sequence variation and human phenotype. Nucleic Acids Res. 2014;42(Database issue):D980–5. doi: 10.1093/nar/gkt1113 gkt1113 [pii]

22. Rath A, Olry A, Dhombres F, Brandt MM, Urbero B, Ayme S. Representation of rare diseases in health information systems: the Orphanet approach to serve a wide range of end users. Hum Mutat. 2012;33(5):803–808. doi:10.1002/humu.22078

23. Pagon RA. GeneReviews. University of Washington; 1993.

24. Peri S, Navarro JD, Kristiansen TZ, et al. Human protein reference database as a discovery resource for proteomics. Nucleic Acids Res. 2004;32(Database issue):D497–501. doi:10.1093/nar/gkh070

25. Geer LY, Marchler-Bauer A, Geer RC, et al. The NCBI BioSystems database. Nucleic Acids Res. 2010;38(Database issue):D492–6. doi:10.1093/nar/gkp858

26. Seal RL, Gordon SM, Lush MJ, Wright MW, Bruford EA. genenames.org: the HGNC resources in 2011. Nucleic Acids Res. 2011;39(Database issue):D514–9. doi:10.1093/nar/gkq892

27. Bovolenta LA, Acencio ML, Lemke N. HTRIdb: an open-access database for experimentally verified human transcriptional regulation interactions. BMC Genomics. 2012;13:405. doi: 10.1186/1471-2164-13-405

28. O’Rawe JA, Wu Y, Dorfel MJ, et al. TAF1 Variants Are Associated with Dysmorphic Features, Intellectual Disability, and Neurological Manifestations. Am J Hum Genet. 2015;97(6):922–932. doi:10.1016/j.ajhg.2015.11.005

29. Swaminathan M, Bannon SA, Routbort M, et al. Hematologic malignancies and Li-Fraumeni syndrome. Cold Spring Harb Mol Case Stud. 2019;5(1). doi:10.1101/mcs.a003210

30. Tanaka AJ, Bai R, Cho MT, et al. De novo mutations in PURA are associated with hypotonia and developmental delay. Cold Spring Harb Mol Case Stud. 2015;1(1):a000356. doi: 10.1101/mcs.a000356

31. Yang H, Douglas G, Monaghan KG, et al. De novo truncating variants in the AHDC1 gene encoding the AT-hook DNA-binding motif-containing protein 1 are associated with intellectual disability and developmental delay. Cold Spring Harb Mol Case Stud. 2015;1(1):a000562. doi:10.1101/mcs.a000562

32. Zimmerman E, Maron JL. FOXP2 gene deletion and infant feeding difficulties: a case report. Cold Spring Harb Mol Case Stud. 2016;2(1):a000547. doi:10.1101/mcs.a000547

33. Tanaka AJ, Cho MT, Retterer K, et al. De novo pathogenic variants in CHAMP1 are associated with global developmental delay, intellectual disability, and dysmorphic facial features. Cold Spring Harb Mol Case Stud. 2016;2(1):a000661. doi:10.1101/mcs.a000661

34. Joshi M, Anselm I, Shi J, et al. Mutations in the substrate binding glycine-rich loop of the mitochondrial processing peptidase-alpha protein (PMPCA) cause a severe mitochondrial disease. Cold Spring Harb Mol Case Stud. 2016;2(3):a000786. doi:10.1101/mcs.a000786

35. Yu HC, Coughlin CR, Geiger EA, et al. Discovery of a potentially deleterious variant in TMEM87B in a patient with a hemizygous 2q13 microdeletion suggests a recessive condition characterized by congenital heart disease and restrictive cardiomyopathy. Cold Spring Harb Mol Case Stud. 2016;2(3):a000844. doi:10.1101/mcs.a000844

36. Leinoe E, Nielsen OJ, Jonson L, Rossing M. Whole-exome sequencing of a patient with severe and complex hemostatic abnormalities reveals a possible contributing frameshift mutation in C3AR1. Cold Spring Harb Mol Case Stud. 2016;2(4):a000828. doi:10.1101/mcs.a000828

37. Griffin LB, Farley FA, Antonellis A, Keegan CE. A novel FGD1 mutation in a family with Aarskog-Scott syndrome and predominant features of congenital joint contractures. Cold Spring Harb Mol Case Stud. 2016;2(4):a000943. doi:10.1101/mcs.a000943

38. Pierce SB, Gulsuner S, Stapleton GA, et al. Infantile onset spinocerebellar ataxia caused by compound heterozygosity for Twinkle mutations and modeling of Twinkle mutations causing recessive disease. Cold Spring Harb Mol Case Stud. 2016;2(4):a001107. doi:10.1101/mcs.a001107

39. Moskowitz AM, Belnap N, Siniard AL, et al. A de novo missense mutation in ZMYND11 is associated with global developmental delay, seizures, and hypotonia. Cold Spring Harb Mol Case Stud. 2016;2(5):a000851. doi:10.1101/mcs.a000851

40. Smedemark-Margulies N, Brownstein CA, Vargas S, et al. A novel de novo mutation in ATP1A3 and childhood-onset schizophrenia. Cold Spring Harb Mol Case Stud. 2016;2(5):a001008. doi:10.1101/mcs.a001008

41. Malcolmson J, Kleyner R, Tegay D, et al. SCN8A mutation in a child presenting with seizures and developmental delays. Cold Spring Harb Mol Case Stud. 2016;2(6):a001073. doi:10.1101/mcs.a001073 MalcolmsonMCS001073 [pii]

42. Kleyner R, Malcolmson J, Tegay D, et al. KBG syndrome involving a single-nucleotide duplication in ANKRD11. Cold Spring Harb Mol Case Stud. 2016;2(6):a001131. doi:10.1101/mcs.a001131

43. Webster E, Cho MT, Alexander N, et al. De novo PHIP-predicted deleterious variants are associated with developmental delay, intellectual disability, obesity, and dysmorphic features. Cold Spring Harb Mol Case Stud. 2016;2(6):a001172. doi:10.1101/mcs.a001172

44. Colby S, Yehia L, Niazi F, et al. Exome sequencing reveals germline gain-of-function EGFR mutation in an adult with Lhermitte-Duclos disease. Cold Spring Harb Mol Case Stud. 2016;2(6):a001230. doi:10.1101/mcs.a001230

45. Yu AC, Chan AY, Au WC, Shen Y, Chan TF, Chan HE. Whole-genome sequencing of two probands with hereditary spastic paraplegia reveals novel splice-donor region variant and known pathogenic variant in SPG11. Cold Spring Harb Mol Case Stud. 2016;2(6):a001248. doi:10.1101/mcs.a001248

46. Polfus LM, Boerwinkle E, Gibbs RA, et al. Whole-exome sequencing reveals an inherited R566X mutation of the epithelial sodium channel beta-subunit in a case of early-onset phenotype of Liddle syndrome. Cold Spring Harb Mol Case Stud. 2016;2(6):a001255. doi:10.1101/mcs.a001255

47. Delpire E, Wolfe L, Flores B, et al. A patient with multisystem dysfunction carries a truncation mutation in human SLC12A2, the gene encoding the Na-K-2Cl cotransporter, NKCC1. Cold Spring Harb Mol Case Stud. 2016;2(6):a001289. doi:10.1101/mcs.a001289

48. Bourne SC, Townsend KN, Shyr C, et al. Optic atrophy, cataracts, lipodystrophy/lipoatrophy, and peripheral neuropathy caused by a de novo OPA3 mutation. Cold Spring Harb Mol Case Stud. 2017;3(1):a001156. doi:10.1101/mcs.a001156

49. Patel RM, Liu D, Gonzaga-Jauregui C, et al. An exome sequencing study of Moebius syndrome including atypical cases reveals an individual with CFEOM3A and a TUBB3 mutation. Cold Spring Harb Mol Case Stud. 2017;3(2):a000984. doi:10.1101/mcs.a000984

50. Morton SU, Prabhu SP, Lidov HGW, et al. AIFM1 mutation presenting with fatal encephalomyopathy and mitochondrial disease in an infant. Cold Spring Harb Mol Case Stud. 2017;3(2):a001560. doi:10.1101/mcs.a001560

51. Caglayan AO, Sezer RG, Kaymakcalan H, et al. ALPK3 gene mutation in a patient with congenital cardiomyopathy and dysmorphic features. Cold Spring Harb Mol Case Stud. 2017;3(5). doi:10.1101/mcs.a001859

52. Inlora J, Sailani MR, Khodadadi H, et al. Identification of a novel mutation in the APTX gene associated with ataxia-oculomotor apraxia. Cold Spring Harb Mol Case Stud. 2017;3(6). doi:10.1101/mcs.a002014

53. Johnston JJ, Lee C, Wentzensen IM, et al. Compound heterozygous alterations in intraflagellar transport protein CLUAP1 in a child with a novel Joubert and oral-facial-digital overlap syndrome. Cold Spring Harb Mol Case Stud. 2017;3(4). doi:10.1101/mcs.a001321

54. Dardour L, Roelens F, Race V, Souche E, Holvoet M, Devriendt K. SPG20 mutation in three siblings with familial hereditary spastic paraplegia. Cold Spring Harb Mol Case Stud. 2017;3(4). doi:10.1101/mcs.a001537

55. Whitford W, Hawkins I, Glamuzina E, et al. Compound heterozygous SLC19A3 mutations further refine the critical promoter region for biotin-thiamine-responsive basal ganglia disease. Cold Spring Harb Mol Case Stud. 2017;3(6). doi:10.1101/mcs.a001909

56. Rohanizadegan M, Abdo SM, O’Donnell-Luria A, et al. Utility of rapid whole-exome sequencing in the diagnosis of Niemann-Pick disease type C presenting with fetal hydrops and acute liver failure. Cold Spring Harb Mol Case Stud. 2017;3(6). doi:10.1101/mcs.a002147

57. Kaiwar C, Zimmermann MT, Ferber MJ, et al. Novel NR2F1 variants likely disrupt DNA binding: molecular modeling in two cases, review of published cases, genotype-phenotype correlation, and phenotypic expansion of the Bosch-Boonstra-Schaaf optic atrophy syndrome. Cold Spring Harb Mol Case Stud. 2017;3(6). doi:10.1101/mcs.a002162

58. Sailani MR, Chappell J, Jingga I, et al. WISP3 mutation associated with pseudorheumatoid dysplasia. Cold Spring Harb Mol Case Stud. 2018;4(1). doi:10.1101/mcs.a001990

59. Tanaka AJ, Cho MT, Willaert R, et al. De novo variants in EBF3 are associated with hypotonia, developmental delay, intellectual disability, and autism. Cold Spring Harb Mol Case Stud. 2017;3(6). doi:10.1101/mcs.a002097

60. Lu JG, Bishop J, Cheyette S, et al. A novel PRRT2 pathogenic variant in a family with paroxysmal kinesigenic dyskinesia and benign familial infantile seizures. Cold Spring Harb Mol Case Stud. 2018;4(1). doi: 10.1101/mcs.a002287

61. Koboldt DC, Mihalic Mosher T, Kelly BJ, et al. A de novo nonsense mutation in ASXL3 shared by siblings with Bainbridge-Ropers syndrome. Cold Spring Harb Mol Case Stud. 2018;4(3). doi:10.1101/mcs.a002410

62. Miller KE, Kelly B, Fitch J, et al. Genome sequencing identifies somatic BRAF duplication c.1794_1796dupTAC;p.Thr599dup in pediatric patient with low-grade ganglioglioma. Cold Spring Harb Mol Case Stud. 2018;4(2). doi:10.1101/mcs.a002618

63. Sanford E, Watkins K, Nahas S, et al. Rapid whole-genome sequencing identifies a novel AIRE variant associated with autoimmune polyendocrine syndrome type 1. Cold Spring Harb Mol Case Stud. 2018;4(3). doi:10.1101/mcs.a002485

64. Berland S, Toft-Bertelsen TL, Aukrust I, et al. A de novo Ser111Thr variant in aquaporin-4 in a patient with intellectual disability, transient signs of brain ischemia, transient cardiac hypertrophy, and progressive gait disturbance. Cold Spring Harb Mol Case Stud. 2018;4(1). doi:10.1101/mcs.a002303

65. Miller CA, Dahiya S, Li T, et al. Resistance-promoting effects of ependymoma treatment revealed through genomic analysis of multiple recurrences in a single patient. Cold Spring Harb Mol Case Stud. 2018;4(2). doi:10.1101/mcs.a002444

66. Bodian DL, Schreiber JM, Vilboux T, Khromykh A, Hauser NS. Mutation in an alternative transcript of CDKL5 in a boy with early-onset seizures. Cold Spring Harb Mol Case Stud. 2018;4(3). doi: 10.1101/mcs.a002360

67. Velez G, Bassuk AG, Schaefer KA, et al. A novel de novo CAPN5 mutation in a patient with inflammatory vitreoretinopathy, hearing loss, and developmental delay. Cold Spring Harb Mol Case Stud. 2018;4(3). doi:10.1101/mcs.a002519

68. Sweeney NM, Nahas SA, Chowdhury S, et al. The case for early use of rapid whole-genome sequencing in management of critically ill infants: late diagnosis of Coffin-Siris syndrome in an infant with left congenital diaphragmatic hernia, congenital heart disease, and recurrent infections. Cold Spring Harb Mol Case Stud. 2018;4(3). doi:10.1101/mcs.a002469

69. Cotter JA, Szymanski L, Karimov C, et al. Transmission of a TP53 germline mutation from unaffected male carrier associated with pediatric glioblastoma in his child and gestational choriocarcinoma in his female partner. Cold Spring Harb Mol Case Stud. 2018;4(2). doi:10.1101/mcs.a002576

70. Antwi P, Hong CS, Duran D, et al. A novel association of campomelic dysplasia and hydrocephalus with an unbalanced chromosomal translocation upstream of SOX9. Cold Spring Harb Mol Case Stud. 2018;4(3). doi:10.1101/mcs.a002766

71. Murry JB, Machini K, Ceyhan-Birsoy O, et al. Reconciling newborn screening and a novel splice variant in BTD associated with partial biotinidase deficiency: a BabySeq Project case report. Cold Spring Harb Mol Case Stud. 2018;4(4). doi:10.1101/mcs.a002873

72. Schwartz JR, Walsh MP, Ma J, et al. Clonal dynamics of donor-derived myelodysplastic syndrome after unrelated hematopoietic cell transplantation for high-risk pediatric B-lymphoblastic leukemia. Cold Spring Harb Mol Case Stud. 2018;4(5). doi:10.1101/mcs.a002980

73. Fomchenko EI, Duran D, Jin SC, et al. De novo MYH9 mutation in congenital scalp hemangioma. Cold Spring Harb Mol Case Stud. 2018;4(4). doi:10.1101/mcs.a002998

74. Grant AR, Hemphill SE, Vincent LM, Rehm HL. Reclassification of the BRAF p.Ile208Val variant by case-level data sharing. Cold Spring Harb Mol Case Stud. 2018;4(5). doi: 10.1101/mcs.a002675

75. Tan QK, Cope H, Spillmann RC, et al. Further evidence for the involvement of EFL1 in a Shwachman-Diamond-like syndrome and expansion of the phenotypic features. Cold Spring Harb Mol Case Stud. 2018;4(5). doi:10.1101/mcs.a003046

76. Koboldt DC, Kastury RD, Waldrop MA, et al. In-frame de novo mutation in BICD2 in two patients with muscular atrophy and arthrogryposis. Cold Spring Harb Mol Case Stud. 2018;4(5). doi:10.1101/mcs.a003160

77. Dubard Gault M, Mandelker D, DeLair D, et al. Germline SDHA mutations in children and adults with cancer. Cold Spring Harb Mol Case Stud. 2018;4(4). doi: 10.1101/mcs.a002584

78. Erdrich J, Schaberg KB, Khodadoust MS, et al. Surgical and molecular characterization of primary and metastatic disease in a neuroendocrine tumor arising in a tailgut cyst. Cold Spring Harb Mol Case Stud. 2018;4(5). doi:10.1101/mcs.a003004

79. Zech M, Lam DD, Weber S, et al. A unique de novo gain-of-function variant in CAMK4 associated with intellectual disability and hyperkinetic movement disorder. Cold Spring Harb Mol Case Stud. 2018;4(6). doi: 10.1101/mcs.a003293

80. Haskell GT, Mori M, Powell C, et al. Combination of exome sequencing and immune testing confirms Aicardi-Goutieres syndrome type 5 in a challenging pediatric neurology case. Cold Spring Harb Mol Case Stud. 2018;4(5). doi:10.1101/mcs.a002758

81. David MP, Venkatramani R, Lopez-Terrada DH, Roy A, Patil N, Fisher KE. Multimodal molecular analysis of an atypical small cell carcinoma of the ovary, hypercalcemic type. Cold Spring Harb Mol Case Stud. 2018;4(5). doi:10.1101/mcs.a002956

82. Khurana M, Edwards D, Rescorla F, et al. Whole-exome sequencing enables correct diagnosis and surgical management of rare inherited childhood anemia. Cold Spring Harb Mol Case Stud. 2018;4(5). doi: 10.1101/mcs.a003152

83. Okur V, Ganapathi M, Wilson A, Chung WK. Biallelic variants in VARS in a family with two siblings with intellectual disability and microcephaly: case report and review of the literature. Cold Spring Harb Mol Case Stud. 2018;4(5). doi:10.1101/mcs.a003301

84. Martignetti JA, Pandya D, Nagarsheth N, et al. Detection of endometrial precancer by a targeted gynecologic cancer liquid biopsy. Cold Spring Harb Mol Case Stud. 2018;4(6). doi:10.1101/mcs.a003269

85. Briggs B, James KN, Chowdhury S, et al. Novel Factor XIII variant identified through whole-genome sequencing in a child with intracranial hemorrhage. Cold Spring Harb Mol Case Stud. 2018;4(6). doi:10.1101/mcs.a003525

86. Tanaka AJ, Okumoto K, Tamura S, et al. A newly identified mutation in the PEX26 gene is associated with a milder form of Zellweger spectrum disorder. Cold Spring Harb Mol Case Stud. 2019;5(1). doi: 10.1101/mcs.a003483

87. Qian Y, Wu B, Lu Y, et al. Early-onset infant epileptic encephalopathy associated with a de novo PPP3CA gene mutation. Cold Spring Harb Mol Case Stud. 2018;4(6). doi: 10.1101/mcs.a002949

88. Sanford E, Farnaes L, Batalov S, et al. Concomitant diagnosis of immune deficiency and Pseudomonas sepsis in a 19 month old with ecthyma gangrenosum by host whole-genome sequencing. Cold Spring Harb Mol Case Stud. 2018;4(6). doi:10.1101/mcs.a003244

89. Claassen D, Boals M, Bowling KM, et al. Complexities of genetic diagnosis illustrated by an atypical case of congenital hypoplastic anemia. Cold Spring Harb Mol Case Stud. 2018;4(6). doi:10.1101/mcs.a003384

90. Windpassinger C, Piard J, Bonnard C, et al. CDK10 Mutations in Humans and Mice Cause Severe Growth Retardation, Spine Malformations, and Developmental Delays. Am J Hum Genet. 2017;101(3):391–403. doi:10.1016/j.ajhg.2017.08.003

91. Lessel D, Schob C, Kury S, et al. De Novo Missense Mutations in DHX30 Impair Global Translation and Cause a Neurodevelopmental Disorder. Am J Hum Genet. 2017;101(5):716–724. doi:10.1016/j.ajhg.2017.09.014

92. Paul A, Drecourt A, Petit F, et al. FDXR Mutations Cause Sensorial Neuropathies and Expand the Spectrum of Mitochondrial Fe-S-Synthesis Diseases. Am J Hum Genet. 2017;101(4):630–637. doi:10.1016/j.ajhg.2017.09.007

93. Watson LM, Bamber E, Schnekenberg RP, et al. Dominant Mutations in GRM1 Cause Spinocerebellar Ataxia Type 44. Am J Hum Genet. 2017;101(3):451–458. doi:10.1016/j.ajhg.2017.08.005

94. Habarou F, Hamel Y, Haack TB, et al. Biallelic Mutations in LIPT2 Cause a Mitochondrial Lipoylation Defect Associated with Severe Neonatal Encephalopathy. Am J Hum Genet. 2017;101(2):283–290. doi:10.1016/j.ajhg.2017.07.001

95. Lake NJ, Webb BD, Stroud DA, et al. Biallelic Mutations in MRPS34 Lead to Instability of the Small Mitoribosomal Subunit and Leigh Syndrome. Am J Hum Genet. 2017;101(2):239–254. doi:10.1016/j.ajhg.2017.07.005

96. Boudin E, de Jong TR, Prickett TCR, et al. Bi-allelic Loss-of-Function Mutations in the NPR-C Receptor Result in Enhanced Growth and Connective Tissue Abnormalities. Am J Hum Genet. 2018;103(2):288–295. doi:10.1016/j.ajhg.2018.06.007

97. Lamers IJC, Reijnders MRF, Venselaar H, et al. Recurrent De Novo Mutations Disturbing the GTP/GDP Binding Pocket of RAB11B Cause Intellectual Disability and a Distinctive Brain Phenotype. Am J Hum Genet. 2017;101(5):824–832. doi:10.1016/j.ajhg.2017.09.015

98. Reijnders MRF, Ansor NM, Kousi M, et al. RAC1 Missense Mutations in Developmental Disorders with Diverse Phenotypes. Am J Hum Genet. 2017;101(3):466–477. doi:10.1016/j.ajhg.2017.08.007

99. Bayram Y, White JJ, Elcioglu N, et al. REST Final-Exon-Truncating Mutations Cause Hereditary Gingival Fibromatosis. Am J Hum Genet. 2017;101(1):149–156. doi:10.1016/j.ajhg.2017.06.006

100. De Mori R, Romani M, D’Arrigo S, et al. Hypomorphic Recessive Variants in SUFU Impair the Sonic Hedgehog Pathway and Cause Joubert Syndrome with Cranio-facial and Skeletal Defects. Am J Hum Genet. 2017;101(4):552–563. doi:10.1016/j.ajhg.2017.08.017

101. Ivanova EL, Mau-Them FT, Riazuddin S, et al. Homozygous Truncating Variants in TBC1D23 Cause Pontocerebellar Hypoplasia and Alter Cortical Development. Am J Hum Genet. 2017;101(3):428–440. doi:10.1016/j.ajhg.2017.07.010

102. Skraban CM, Wells CF, Markose P, et al. WDR26 Haploinsufficiency Causes a Recognizable Syndrome of Intellectual Disability, Seizures, Abnormal Gait, and Distinctive Facial Features. Am J Hum Genet. 2017;101(1):139–148. doi:10.1016/j.ajhg.2017.06.002

103. Guella I, McKenzie MB, Evans DM, et al. De Novo Mutations in YWHAG Cause Early-Onset Epilepsy. Am J Hum Genet. 2017;101(2):300–310. doi:10.1016/j.ajhg.2017.07.004

104. Wang K, Li M, Hakonarson H. ANNOVAR: functional annotation of genetic variants from high-throughput sequencing data. Nucleic Acids Res. 2010;38(16):e164. doi:10.1093/nar/gkq603

105. Zeng S, Zhang M, Wang X, et al. Long-read sequencing identified intronic repeat expansions in SAMD12 from Chinese pedigrees affected with familial cortical myoclonic tremor with epilepsy. J Med Genet. 2019;56(4):265–270. doi:10.1136/jmedgenet-2018-105484

106. Ishiura H, Doi K, Mitsui J, et al. Expansions of intronic TTTCA and TTTTA repeats in benign adult familial myoclonic epilepsy. Nat Genet. 2018;50(4):581–590. doi:10.1038/s41588-018-0067-2

107. Chaisson MJP, Sanders AD, Zhao X, et al. Multi-platform discovery of haplotype-resolved structural variation in human genomes. Nat Commun. 2019;10(1): 1–16. doi:10.1038/s41467-018-08148-z

108. Zook JM, Hansen NF, Olson ND, et al. A robust benchmark for germline structural variant detection. bioRxiv. July 2019:664623. doi:10.1101/664623

109. Mehrabi S, Krishnan A, Sohn S, et al. DEEPEN: A negation detection system for clinical text incorporating dependency relation into NegEx. J Biomed Inform. 2015;54:213–219. doi:10.1016/j.jbi.2015.02.010

110. Chapman WW, Bridewell W, Hanbury P, Cooper GF, Buchanan BG. A simple algorithm for identifying negated findings and diseases in discharge summaries. J Biomed Inform. 2001;34(5):301–310. doi:10.1006/jbin.2001.1029

111. Kircher M, Witten DM, Jain P, O’Roak BJ, Cooper GM, Shendure J. A general framework for estimating the relative pathogenicity of human genetic variants. Nat Genet. 2014;46(3):310–315. doi:10.1038/ng.2892

112. Ioannidis NM, Rothstein JH, Pejaver V, et al. REVEL: An Ensemble Method for Predicting the Pathogenicity of Rare Missense Variants. Am J Hum Genet. 2016;99(4):877–885. doi:10.1016/j.ajhg.2016.08.016

113. Havrilla JM, Pedersen BS, Layer RM, Quinlan AR. A map of constrained coding regions in the human genome. Nat Genet. 2019;51(1):88–95. doi:10.1038/s41588-018-0294-6

