## Supplemental Materials and Methods for "Phen2Gene: Rapid Phenotype-Driven Gene Prioritization for Rare Diseases"

### Supplementary Materials

**Supplemental Material 1. HPO terms for all benchmark sets.** This file contains the HPO terms for each patient for each benchmark dataset.

https://github.com/WGLab/Phen2Gene/releases/download/1.1.0/testing_data.zip

| Case | Causal Gene | Phenotype Examples |
|---|---|---|
| 1 | SNAP25 | profound static encephalopathy, generalized epilepsy |
| 2 | COL10A1 | skeletal dysplasia, developmental delay |
| 3 | ARID1B | cystic hygroma, aryngomalacia |
| 4 | SCN1A | generalized seizures, developmental delay |
| 5 | CDKL5 | Seizure, hypotonia, global developmental delay |
| 6 | MYH10 | Microcephaly, hydrocephalus |
| 7 | LMNA | atrial fibrillation, hypertrophic cardiomyopathy |
| 8 | ALG13 | Encephalopathy, infantile spasms |
| 9 | EHMT1 | developmental delay, hypotonia |
| 10 | SLC1A4 | Congenital microcephaly, developmental delay, eczema |
| 11 | MAN2B1 | dilated cardiomyopathy, developmental delay, hypotonia |
| 12 | MYO7A | hearing loss, retinitis pigmentosa, visual loss |
| 13 | ARID1B | profound static encephalopathy, generalized epilepsy |
| 14 | EHMT1 | central hypothyroidism, laryngomalacia |
| 15 | PTEN | pulmonic stenosis, polydactyly |
| 16 | ATRX | Dwarfism, macrocephaly |
| 17 | TKT | dysmorphic features, undescended testis |
| 18 | PLA2G4A | failure to thrive, developmental delay |
| 19 | DDX3X | hypertrophic cardiomyopathy, Noonan syndrome |
| 20 | HNRNPH2 | pervasive developmental disorder, global developmental delay |
| 21 | PTPN11 | global developmental delay, failure to thrive |
| 22 | COL7A1 | Cardiomyopathy, neonatal respiratory distress |
| 23 | KMT2D | dystrophic epidermolysis bullosa, Abnormal blistering of the skin |
| 24 | SHH | cleft palate, Unilateral renal dysplasia |
| 25 | NAA15 | Microcephaly, Toe clinodactyly |
| 26 | CDKL5 | heterotaxy syndrome, congenital heart disease |
| 27 | POMT1 | developmental delay, seizures |

**Supplemental Table 1. 27 Expert Curation Cases from Son et al.**

| Case | PubMed ID | Causal Gene |
|---|---|---|
| 1 | 27148565 | PURA |
| 2 | 27148574 | AHDC1 |
| 3 | 27148578 | FOXP2 |
| 4 | 27148580 | CHAMP1 |
| 5 | 27148589 | PMPCA |
| 6 | 27148590 | TMEM87B |
| 7 | 27551680 | C3AR1 |
| 8 | 27551683 | FGD1 |
| 9 | 27551684 | C10ORF2 |
| 10 | 27626064 | ZMYND11 |
| 11 | 27626066 | ATP1A3 |
| 12 | 27900360 | SCN8A |
| 13 | 27900361 | ANKRD11 |
| 14 | 27900362 | PHIP |
| 15 | 27900366 | EGFR |
| 16 | 27900367 | SPG11 |
| 17 | 27900368 | SCNN1B |
| 18 | 27900370 | SLC12A2 |
| 19 | 28050599 | OPA3 |
| 20 | 28299356 | TUBB3 |
| 21 | 28299359 | AIFM1 |
| 22 | 28630369 | ALPK3 |
| 23 | 28652255 | APTX |
| 24 | 28679688 | CLUAP1 |
| 25 | 28679690 | SPG20 |
| 26 | 28679690 | SPG20 |
| 27 | 28679690 | SPG20 |
| 28 | 28696212 | SLC19A3 |
| 29 | 28802248 | NPC1 |
| 30 | 28963436 | NR2F1 |
| 31 | 28963436 | NR2F1 |
| 32 | 29092958 | WISP3 |
| 33 | 29162653 | EBF3 |
| 34 | 29167286 | PRRT2 |
| 35 | 29305346 | ASXL3 |
| 36 | 29434027 | BRAF |
| 37 | 29437776 | AIRE |
| 38 | 29437797 | AQP4 |
| 39 | 29440180 | MEN1 |
| 40 | 29444904 | GABRA1 |
| 41 | 29472286 | CAPN5 |
| 42 | 29549119 | ARID1B |

| | | |
|---|---|---|
| 43 | 29581140 | TP53 |
| 44 | 29695406 | SOX9 |
| 45 | 29728376 | BTD |
| 46 | 29891567 | ASXL1 |
| 47 | 29903892 | MYH9 |
| 48 | 29945942 | BRAF |
| 49 | 29970384 | EFL1 |
| 50 | 30054298 | BICD2 |
| 51 | 30054298 | BICD2 |
| 52 | 30068732 | SDHA |
| 53 | 30087100 | BCRA1 |
| 54 | 30262571 | CAMK4 |
| 55 | 30275001 | SAMHD1 |
| 56 | 30275002 | SMARCA4 |
| 57 | 30275003 | SPTA1 |
| 58 | 30275004 | VARS |
| 59 | 30275004 | VARS |
| 60 | 30301868 | PTEN |
| 61 | 30404926 | F13A1 |
| 62 | 30446579 | PEX26 |
| 63 | 30455226 | PPP3CA |
| 64 | 30559311 | BTK |
| 65 | 30559313 | CECR1 |
| 66 | 30709875 | TP53 |
| 67 | 30709875 | TP53 |
| 68 | 30709875 | TP53 |
| 69 | 30709875 | TP53 |
| 70 | 30709875 | TP53 |
| 71 | 30709875 | TP53 |
| 72 | 30709875 | TP53 |

**Supplemental Table 2. 72 cases from Cold Spring Harbor Molecular Case Study**

| PubMed ID | Causal Gene | Number of Cases |
|---|---|---|
| 28886341 | CDK10 | 8 |
| 29100085 | DHX30 | 12 |
| 28965846 | FDXR | 8 |
| 28886343 | GRM1 | 4 |
| 28757203 | LIPT2 | 3 |
| 28777931 | MRPS34 | 6 |
| 30032985 | NPR3 | 3 |
| 29106825 | RAB11B | 5 |
| 28886345 | RAC1 | 7 |
| 28686854 | REST | 3 |
| 28965847 | SUFU | 2 |
| 28823707 | TBC1D23 | 3 |
| 28686853 | WDR26 | 13 |
| 28777935 | YWHAG | 6 |

**Supplemental Table 3. 83 Cases from American Journal of Human Genetics**

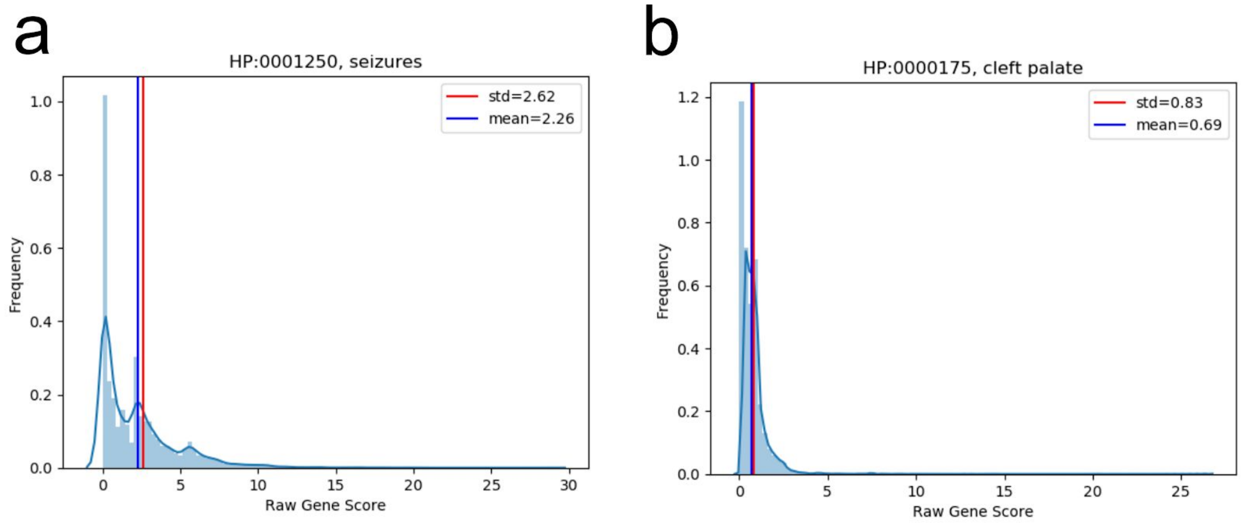


**Supplemental Figure 1. Distribution of Terms Cleft Palate and Seizures.** Two HPO terms (HP:0001250, “seizures” and HP:0000175, “cleft palate”) and their distribution of Enhanced Phenolyzer scores for all linked genes. “Seizures” has less of a negative skew than “cleft palate” and is a less specific HPO term. (a) “Seizures” gene score distribution. (b) “Cleft palate” gene score distribution.

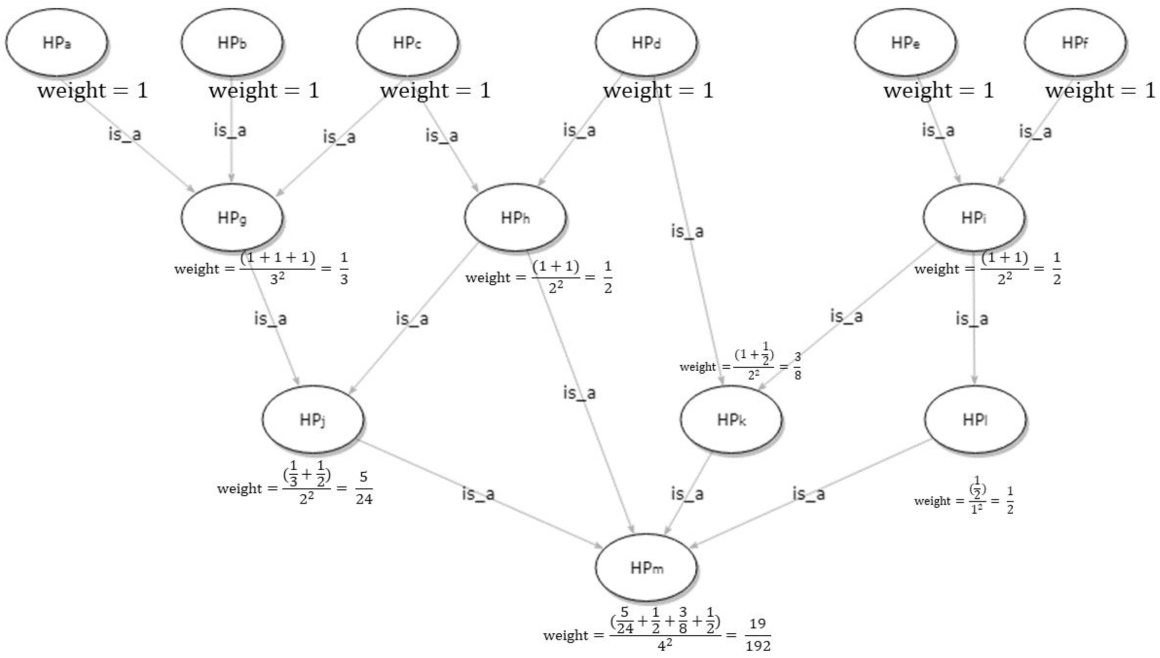


**Supplemental Figure 2. Example of Weight Assignment Calculation.** This figure demonstrates how leaf depth affects the score for an HPO term.

### Supplemental Methods

#### Gene score computation with unweighted HPO terms

Given a set of HPO terms, $TermSet = \left\{HP_j\right\}$ , Phen2Gene searches candidate genes in each $HP_j$'s candidate gene list. Each candidate gene gets one score when it appears in one $HP_j$'s candidate gene list. Then Phen2Gene sums up scores for each candidate gene,

$$S_{unweighted}(gene_i) = \sum_j \delta(gene_i, HP_j) \quad , HP_j \in \left\{HP_j\right\} ,$$

where $S(gene_i, HP_j)$ is $gene_i$'s score in $HP_j$'s candidate gene list. $S\left(gene_i, HP_j\right) = 0$ , if $gene_i$ is not a candidate gene of $HP_j$ . All of genes are sorted by their scores in descending order. All of the candidate genes are sorted by their scores in descending order. If multiple candidate genes receive the same score, with the same rankings, Phen2Gene will give them an average rank. For example, if 100 genes tie at the highest score, their actual rankings are averaged to be 50, rather than 1.

##### Weighting by information content

Child terms in the HPO tree provide more concrete phenotypic information than parent terms, and thus intuitively they should have greater weight than parent terms. "Phenotypic abnormality" (HP:0000118) had 9,399 descendent terms that have no child terms. Generally, these possess the most detailed phenotypic information, and so we assigned the same initial weight of 1 to them. Then, we used the following formula (option 'w' for weight),

$$weight\,(parent\ term) = \frac{\sum_i weight(child\ term_i)}{(number\ of\ children\ terms)^2}$$

to assign the weights for the parental and ancestral terms (**Supplemental Figure 2**).

The user can also assign weights based on ontology-based information content computation[1]. This was not used in the manuscript figures either (option 'ic' for weight), but it is an option that considers different concreteness among leaf terms in the HPO ontology tree,

$$weight\,(HP_i) = -log\left(\frac{\frac{|leaf(HP_i)|+1}{|ancestor(HP_i)|}}{|leaf(HP_j)|+1}\right),$$

where $|leaf\,(HP_i)|$ is the number of leaf descendants $HP_i$ has, $|ancestor\,(HP_i)|$ is the number of ancestors up to "Phenotypic abnormality" (HP:0000118) that $HP_i$ has. This approach also distinguishes leaf terms in HPO tree by the number of ancestral terms those leaf terms have. Finally, the weights are normalized from 0 to 1.

### References


1. Sánchez, D., Batet, M. & Isern, D. Ontology-based information content computation. *Knowl.-Based Syst.* **24**, 297–303 (2011).
